# New Insights into the Conformational Activation of Full-Length Integrin

**DOI:** 10.1101/203661

**Authors:** Tamara C. Bidone, Anirban Polley, Aleksander Durumeric, Tristan Driscoll, Daniel Iwamoto, David Calderwood, Martin A. Schwartz, Gregory A Voth

## Abstract

Integrin binding to extracellular matrix proteins is regulated by conformational transitions from closed, low affinity states to open, high affinity states. However, the pathways of integrin conformational activation remain incompletely understood. Here, by combining all-atom molecular dynamics simulation, coarse-graining, heterogeneous elastic network modeling, and experimental ligand binding measurements, we test the effect of integrin β mutations that destabilize the closed conformation. Our results support a “deadbolt” model of integrin activation, where extension of the headpiece is not coupled to leg separation, consistent with recent cryo-EM reconstructions of integrin intermediates. Moreover, our results are inconsistent with a “switchblade-like” mechanism. The data show that locally correlated atomistic motions are likely responsible for extension of integrin headpiece before separation of transmembrane legs, without persistence of these correlations across the entire protein. By combining modeling and simulation with experiment, this study provides new insight into the structural basis of full-length integrin activation.

## INTRODUCTION

Integrins are transmembrane receptors that signal bidirectionally across the plasma membrane to regulate cell processes such as adhesion, migration, differentiation, and mechanosensing (1–6). Integrins can be found in either bent conformations that bind to extracellular matrix (ECM) ligands with low affinity or in open, high affinity conformations (6–9). The structure of integrin resembles a large head and two legs, with the head containing sites for ligand binding. Both head and legs comprise several subdomains, interconnected by linkers. Owing to the large dimensions of the receptor and the complex interconnections between and within subdomains, that determine activation state, how integrin transits from bent to extended conformations is not totally understood.

Integrins are heterodimers composed of an α and a β subunit that associate non-covalently. For integrin αvβ3, the two subunits form a structure with an extracellular ligand-binding headpiece, two transmembrane helices and two short cytoplasmic tails (10) (Figure 1). The αv subunit consists of five extracellular domains: a seven-bladed β-*propeller* and a *thigh* domain, which is connected by the flexible *linker 1* to the *calf* domains, followed by the *transmembrane* and *cytoplasmic* or *tail* domains. The β3 subunit has seven domains with flexible interconnections: the *β*A *domain* is inserted into the *hybrid* domain, which is, in turn, inserted in the *plexin-semaphorin-integrin* (PSI) domain; these domains are followed by four cysteine-rich *epidermal growth factor* (EGF) modules and the *cytoplasmic* domain (11). The structure of the bent, low affinity integrin αIIbβ3, which is closely related to αvβ3, is represented in Figure 1a.

**Figure 1.**
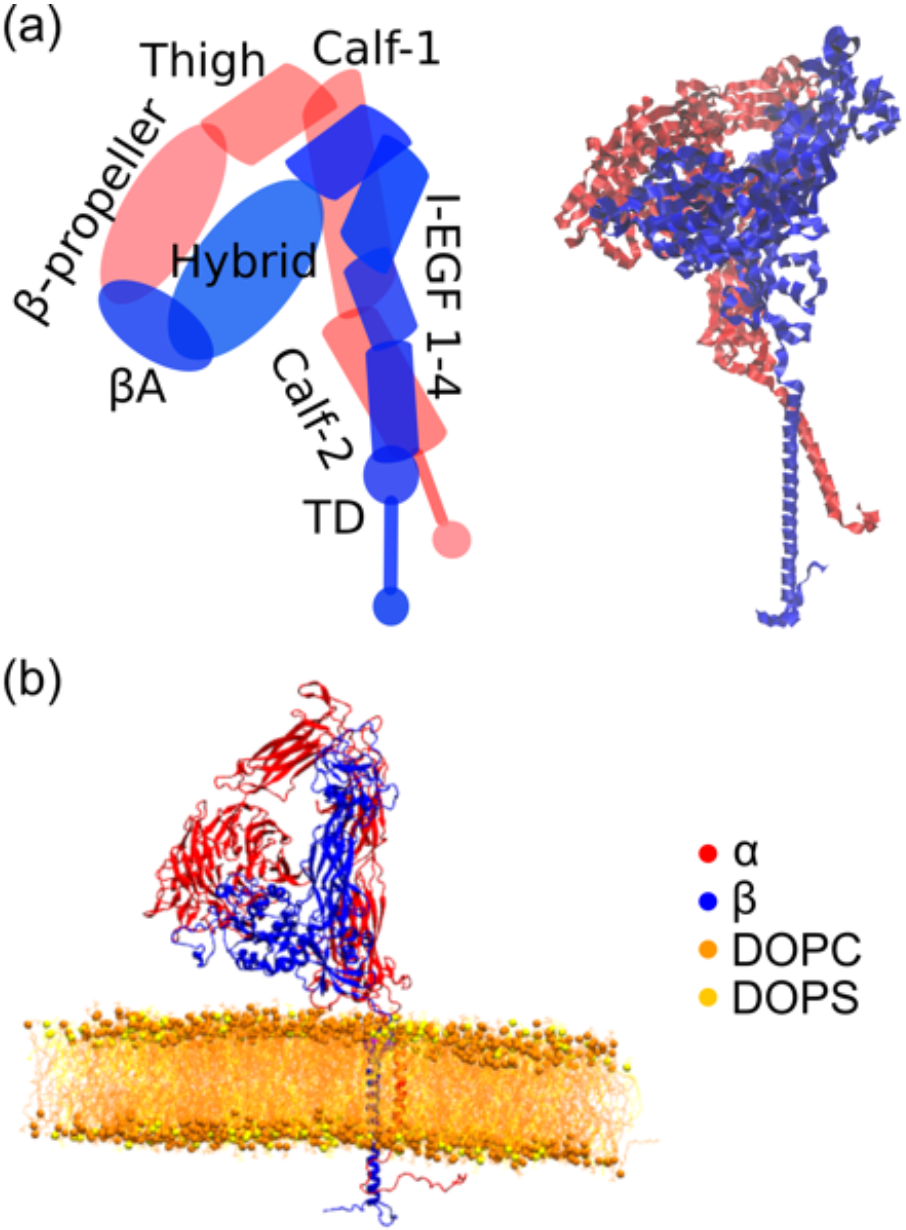
Structures of full length, closed α_IIb_β_3_ and α_V_β_3_ integrins and corresponding molecular models. (a) Structure of α_IIb_β_3_ integrin in its closed, low affinity conformation. Subunits α and β are represented in red and blue, respectively. Corresponding ribbon representation of closed α_IIb_β_3_ integrin on the right (from (24)). (b) Ribbon representation of atomistic full-length α_V_β_3_ integrin embedded in DOPC/DOPS (4:1) lipid bilayer.

Integrin activation occurs via a large scale conformational rearrangement involving relative motions between the α and β subunits and their subdomains on the order of nanometers. Previous studies showed that localized changes of the βA/hybrid interface of the β subunit of both αIIbβ3 and αvβ3 integrins, are critical for these movements (12–14). For example, the α7 helix of the βA domain shifts towards the hybrid domain in a piston-like movement that causes the hybrid domain to swing out by ~60° (8, 15, 16). Also, upon activation, Glu318 in the α subunit β-*propeller* domain becomes bound to the β subunit βA metal site (15, 17). These localized rearrangements in the integrin headpiece shift the conformation from bent to open (8, 12–14). Molecular dynamics (MD) simulations on αvβ3, in combination with steered molecular dynamics (SMD) studies, have suggested that inter-domain contacts between the legs and headpiece raise the energy barrier that must be overcome for opening the βA/hybrid hinge (18). Therefore, concomitantly with the reorganization of both α and β subunits in the integrin headpiece, opening of αvβ3 involves breaking the extensive interfaces between the headpiece and lower legs, and this is valid also for α_IIb_β_3_ (12).

Integrin activation also requires separation of the transmembrane legs. Previous work has studied the initiation of integrin opening by focusing on segments of the integrin headpiece at the nanosecond timescale (19). Less is known about how headpiece motions coordinate with leg separation, which occurs on the second timescale. In fact, several studies of integrin activation examined proteins with mutated, truncated, or entirely absent legs, thus preventing this kind of analysis (16, 20–22).

In order to explain the intrinsically multiscale mechanism of integrin opening, two conceptual models based on experimental observations have been proposed. In the “switchblade” model (8), opening of the βA/hybrid hinge and separation of transmembrane legs occur in a coordinated fashion. In contrast, the “deadbolt” model proposes more conservative changes around the bent structure with progressive loss of constraining contacts between βA domain and β tails that occur before leg separation (23). Thus, without dynamic, nonequilibrium information about how structural changes in the headpiece are coupled to the separation of the legs, major questions remain concerning the pathway of integrin opening.

Recently, structures captured in various degrees of opening of full-length α_IIb_β_3_ integrin showed headpiece extension without leg separation (24). However, even in this scenario, it is still possible that electrostatic interactions in the β_3_ helix decrease upon headpiece extension and that a coordinated structural change occurs at the interface between the two legs without detectable separation, consistent with coordinated structural reconfiguration between headpiece and legs at short length and time scales.

In order to understand the relationship between headpiece extension and legs separation, we have combined here a multiscale simulation approach with experimental ligand binding measurements. We tested the effect of activating mutants on the molecular structure and their impact on the long-range structural rearrangements of integrin. Our results support the notion that headpiece extension occurs before legs separation, consistent with a deadbolt model of integrin activation and inconsistent with a switchblade model. This mechanism is mediated by local, correlated atomistic motions within and between neighboring subdomains of the receptor, and independent from long-range interactions.

## RESULTS

### All atom molecular dynamics simulations of integrin mutants

Inspections of root mean square deviation (RMSD) plots of WT, single, and double integrin mutants showed that all structures reached local equilibrium states within 1 μs of MD simulations (Figure 2a,d). Analysis of root mean square (RMS) fluctuations also showed that some of the most flexible regions of integrin are at the interface between the β-propeller and βA domains, together with the Linker 1 and EGF motifs, which form the α and β genu, respectively (Figure 2b-c, e-f). There was no significant difference in RMS fluctuations between WT and the mutant integrins considered here. These data collectively support that the EGF domain region is relatively plastic, especially between EGF1 and EGF2, at the β knee, and at the PSI/hybrid and hybrid/I-EGF1 junctions. It was previously reported that the flexibility of the βA domain would also facilitate such interdomain interactions (15).

**Figure 2.**
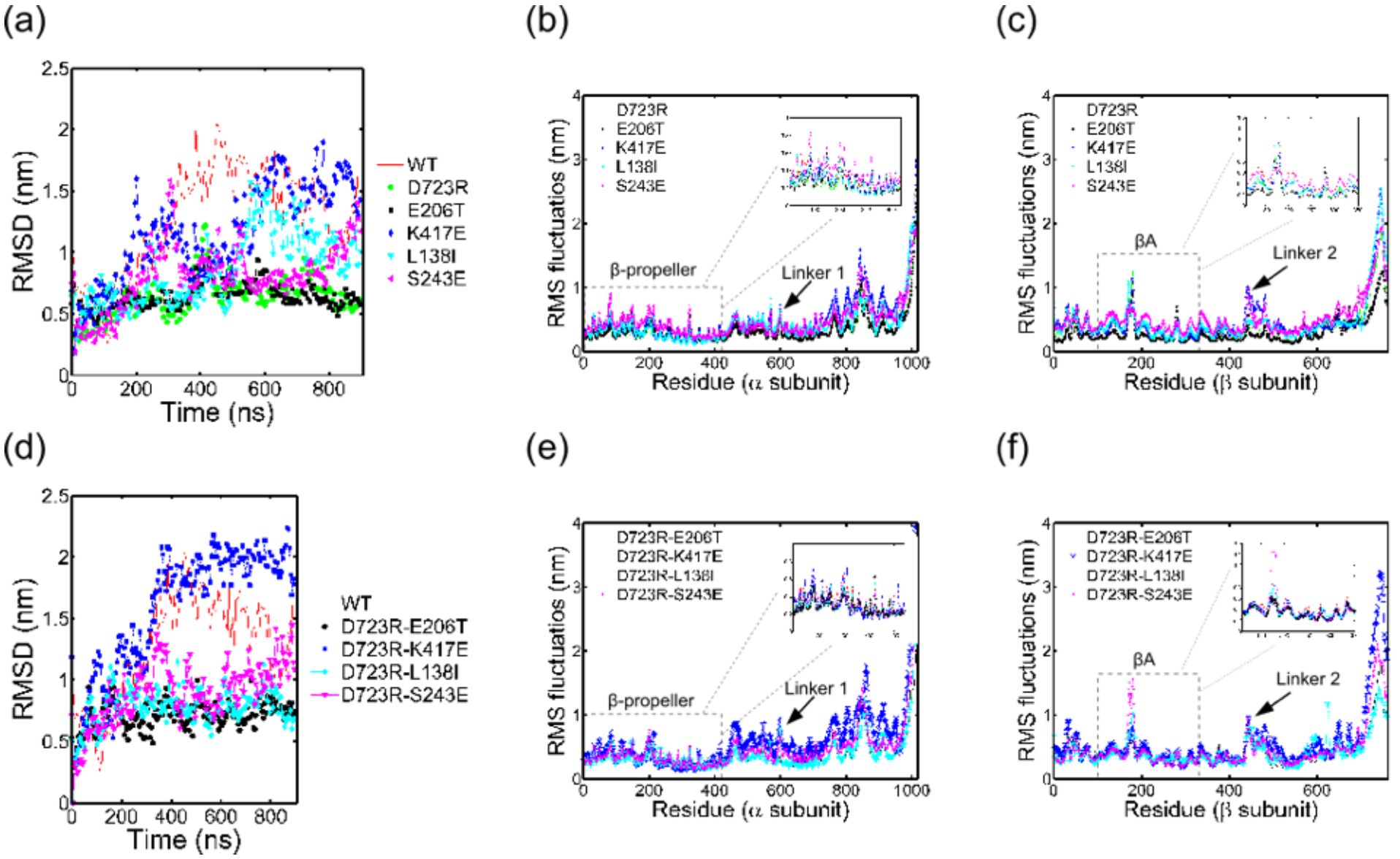
All atom molecular dynamics simulations of integrin αvβ3 mutants. (a) Time evolution of root mean square deviations of the atomistic C_α_ for WT and single integrin mutants relative to the corresponding equilibrated configurations used as input to the MD. (b) Root mean square fluctuations of individual residues in the α subunit of WT and single integrin mutants. (c) Root mean square fluctuations of individual residues in the α subunit of WT and single integrin mutants. Color code in (a), (b), and (c) is: WT (red), D723R (green), E206T (black), K417E (blue), L138I (cyan), S243E (magenta). (d) Time evolution of root mean square deviations of the atomistic C_α_ for WT and double integrin mutants relative to the corresponding equilibrated configurations given as input to the MD. (e) Root mean square fluctuations of individual residues in the α subunit of WT and double integrin mutants. (f) Root mean square fluctuations of individual residues in the α subunit of WT and double integrin mutants. Color code in (d), (e), and (f) is: WT (red), D723R-E206T (black), D723R-K417E (blue), D723R-L138I (cyan), D723R-S243E (magenta).

Over the last 100 ns of the AA MD simulations, the average angle < ϑ > of the mutants was about 5-7% different from WT (Figure 3b); D_12_ was enhanced in the S243E mutant relative to both WT and other single/double mutants (figure 3c); D_EM_ was about 15% different for the single mutants and about 8% different in the double mutants relative to WT (Figure 3d). Accordingly, the single mutants showed enhanced persistency of high values of D_EM_ (Figure 3e). Time evolution of D_12_, D_EM_, and corresponding probability distributions, are reported in Figure S1 and FigureS2 of the Supplemental Information. Time evolutions of ϑ_1_ and ϑ_2_ are reported in Figure S3. Both maximum values and standard deviations of ϑ_1_ and ϑ_2_ over the 1μs-long MD simulations were higher in S243E mutant than the other integrin single/double mutants (Figure S4a-b). Maximum value and standard deviation of ϑ_1_, over 1μs-long MD, were among the highest for D723R with respect to the other single mutants (Figure S4a-b). High standard deviations in the kink angles of S243E and D723R with respect to the other single mutants indicate that the configurations of the two headpiece hinges were far from their mean. Thus, the angles were more flexible in these mutant relative to the other single mutants examined. Also, maximum value and standard deviation for D_12_ were higher in the S243E and D723R (Figures S4c) relative to the other single mutants. Among the double mutants, D723R-E206T also showed high values of peak and standard deviation for kink angle, leg separation, and headpiece extension (Figure S4a-d). Taken together, these results support that the mutants here tested destabilize the integrin closed conformation by enhancing variability of kink angles in both subunits, leg separation, and/or overall distance of the headpiece from the transmembrane legs.

**Figure 3.**
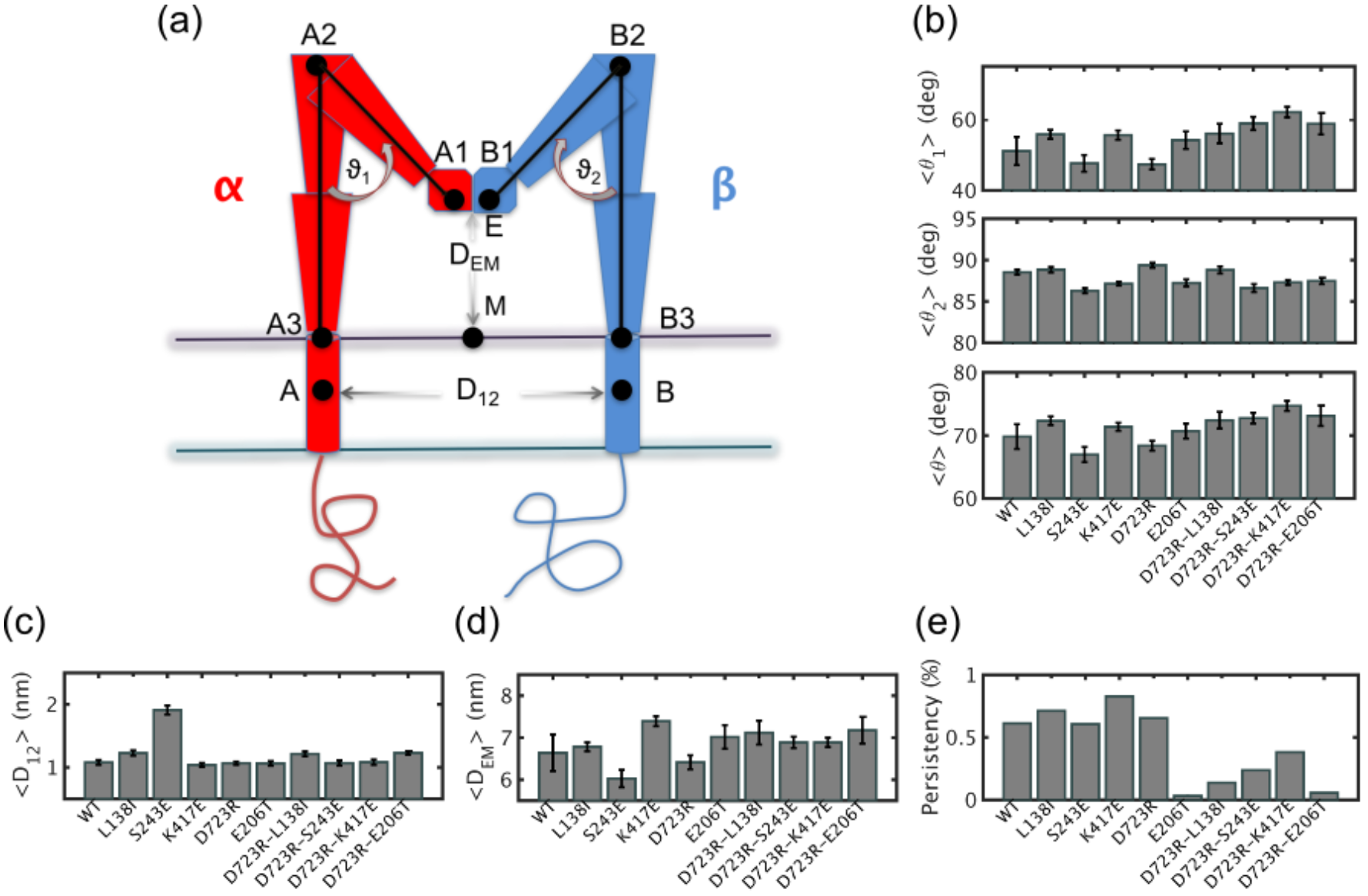
Leg separation, headpiece extension and kink angles in the last 100 ns of AA MD simulations. (a) Schematic diagram of αvβ3 in a lipid bilayer membrane, with red and blue elements representing α and β subunits, respectively. The horizontal lines indicate upper and lower membrane leaflets. Points A1, A2, A3 on the α subunit and points B1, B2, B3 on the β subunit are used to characterize corresponding kink angles, ϑ_1_ and ϑ_2_. Distance D_12_ indicates separation between the two transmembrane helices and D_EM_ is a measure of headpiece extension from the membrane. (b) Average values of ϑ_1_, ϑ_2_ and ϑ in WT and mutant integrins, computed between 900-1000 ns of MD simulations. (c) Average values of D_12_ in WT and mutant integrins. (d) Average values of D_EM_ in WT and mutant integrins. (e) Persistence of extended state for WT and mutant integrins, computed as the fraction of time, between 900-1000nm of MD, that D_EM_ was at least 95% of its maximum value

We next addressed the molecular mechanism by which S243E, one of the most activating mutant, affects integrin conformation. In WT integrin, neutral histidine 244, negatively charged aspartic acid 113, and positively charged arginine 352 surround S243, with Arg352 at a distance (Figure 4a). In MD simulations of systems with protonated glutamic acid in S243E, a salt-bridge formed between the oxygen atom of arginine and the hydrogen atom of glutamic acid (Figure 4b). Salt bridge formation was accompanied by reorientation of the positively charged arginine, which moved closer to the negative charged glutamic acid and re-oriented towards the negatively charged aspartic acid (figure 4b). These molecular rearrangements resulted in greater headpiece extension, flattening of kink angles and separation of the transmembrane legs in the MD simulations. Time evolution and probability density functions of distances of ARG352 from the center of mass of residues 244, 243 and 113 in WT and mutant conditions are in Figure S5.

**Figure 4.**
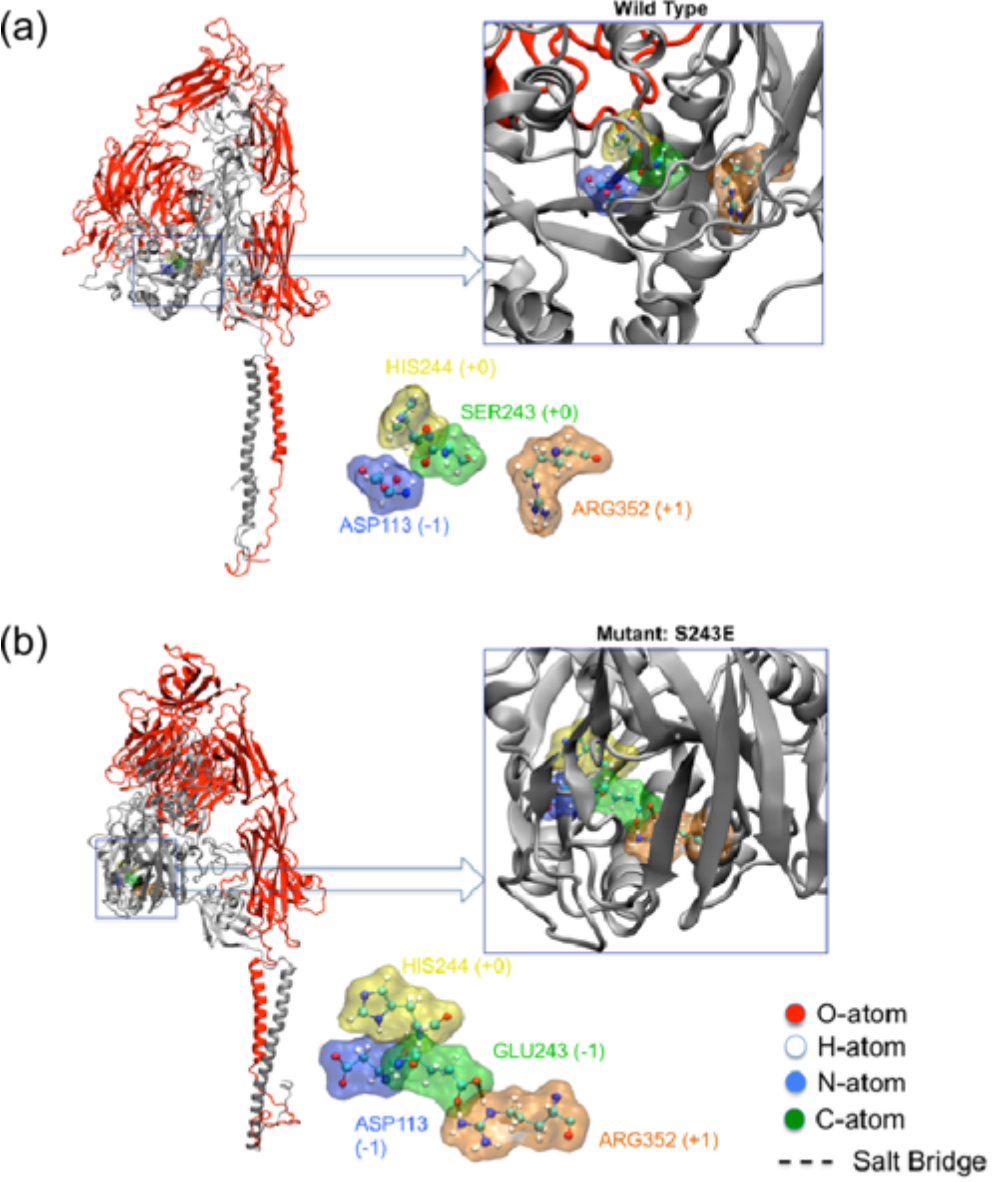
S243E triggers formation of a salt bridge in the βA domain of β subunit. (a) Cartoon representation of full length WT integrin with α subunit in red and β subunit in grey. Highlighted and zoomed are the amino acids surrounding S243 (green): histidine (yellow), aspartic acid (violet) and arginine (orange) (b). Cartoon representation of full length mutant integrin with α subunit in red and β subunit in grey. Highlighted and zoomed are the amino acids surrounding mutated S243E, using the same color code as in (a). Dashed line shows a salt bridge between negative Glu243 and positive Arg352. Oxygen, hydrogen, nitrogen and carbon atoms are shown in ball and stick representation in red, white, blue and green, respectively.

Taken together, the analysis of MD simulations showed local destabilization of the closed configuration in integrin mutants. However, structural quantities which are representative of integrin conformational activation at the level of the whole receptor, such as D_EM_, ϑ_1_ and ϑ_2_, did not convergence within 1 μs. In order to allow for enhanced sampling of integrin long-range interactions, we instead used the MD trajectories to build CG models.

### Coarse-grained (CG) simulations

Atomistic integrin structures were converted into CG systems, without the lipid bilayer included explicitly in the CG model, to detect large-scale motions of the receptor. As described earlier, we used a combination of ED-CG and heteroENM to create the CG model and systematically removed harmonic bonds or converted them into Morse potentials, motivated by the assumption that weak CG effective harmonic correlations between domains are likely to dissociate upon integrin activation. We analyzed our results to identify structural differences between WT and mutant integrins. Our goal was to sample multiple integrin states underlying the equilibrium conformers and to provide insight concerning which mutants most potently destabilize integrin closed states. We mapped the atomistic WT system to a CG ED-CG model of 200 CG sites or “beads”, with average resolution 8 **±** 3 residues per CG site, which is of the same order used previously (32). We compared the CG root mean square fluctuations from ED-CG-heteroENM at different initial cutoffs, spanning 3-5 nm, with those from atomistic simulations converted into CG fluctuations (Figure 5a). Using cutoffs of 3, 4, or 5 nm, the average differences in RMS fluctuations from the all-atom fluctuations were 0.11, 0.12 and 0.16 nm, respectively. We therefore chose a 3 nm cutoff for our CG systems that best reproduced atomistic fluctuations, and built ED-CG-heteroENM models for each mutant integrin (Figure 5b). The fraction of intra-domain springs was about 0.6 in all systems, with inter-domain springs that connected non-consecutive subdomains along the primary sequence below 0.2. Intra-domain connections had the highest spring constants, up to 25 kcalmol^−1^A^2^ (Figure 5d), while inter-domain springs had about 3-fold lower characteristic spring constants, below 8 kcalmol^−1^A^2^ (Figure 5e-f). Snapshots from a representative CG simulation showed average kink angles between 120-140 deg (Figure 6a), depending on the particular mutant. Theses angles are about twice those from MD, showing that conformations far from equilibrium were sampled with this CG method and that the structures are extended. All mutants showed higher kink angles than WT αvβ3 (Figure 6a). Also, comparison of the fraction of time that D_EM_ is above 95% of its maximum showed that all the mutants were above 15%, whereas WT integrin was below 10% (Figure 6b). Thus, in the CG simulations, the mutants were in extended conformations more often than WT. By capturing the effect of point mutations, our CG models of integrin were able to enhance sampling conformations from MD trajectories.

**Figure 5.**
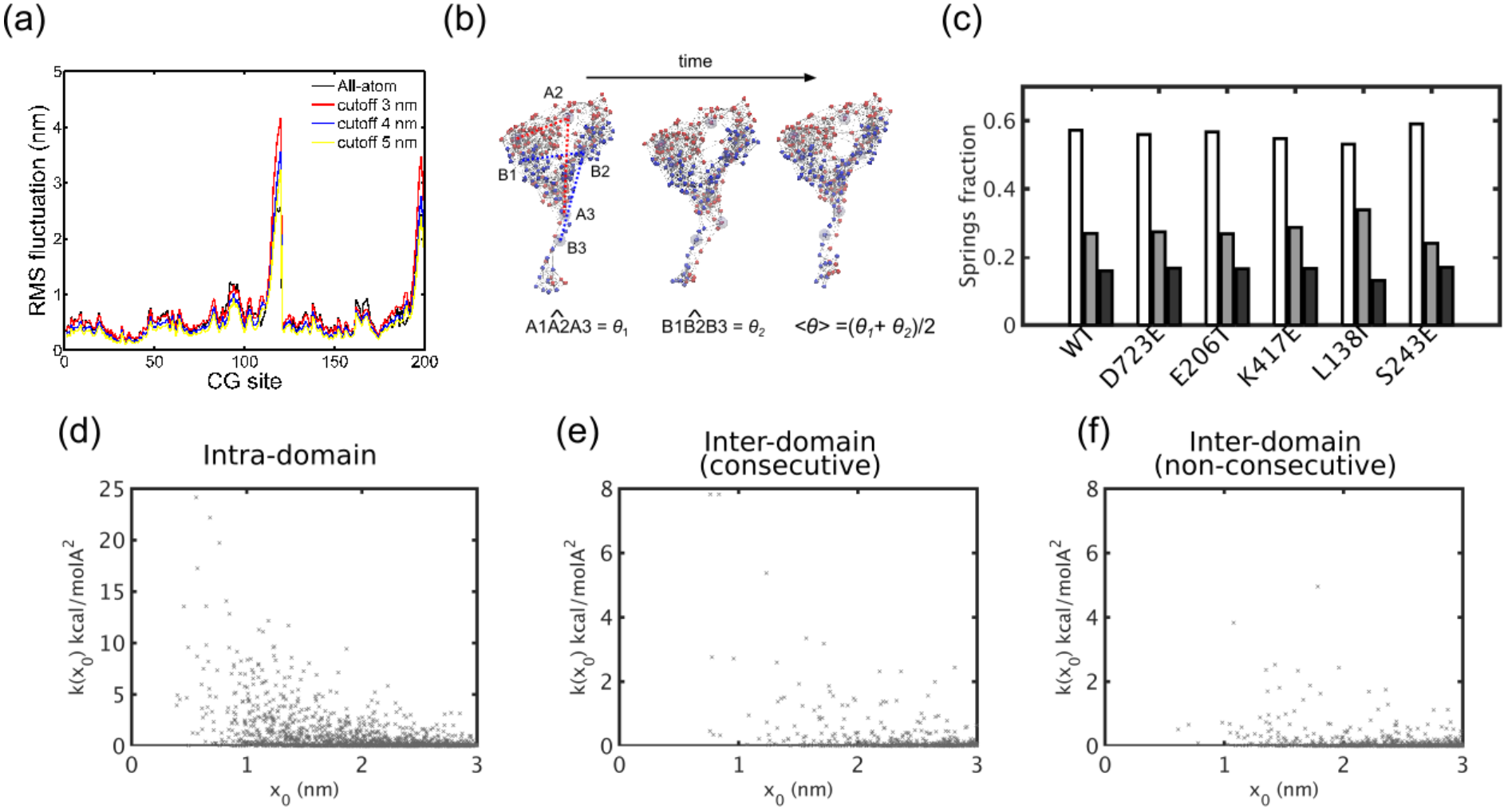
ED-CG-heteroENM models of integrin mutants. (a) Per-CG-site root mean square fluctuations of WT integrin computed from AA MD simulations and ED-CG-heteroENM integrin using cutoffs of 3, 4, and 5 nm. (b) Snapshots from CG-heteroENM S243E integrin simulations, with red indicating CG-sites of the β subunit and blue indicating CG-sites of the β subunit. Bonds represent harmonic interactions. CG-sites representing A1, A2, A3 and B1, B2, B3 are mapped from atomistic residues. (c) Fraction of harmonic interactions from hetero-ENM, for WT and single mutant integrins: interactions within each domain (white); between consecutive subdomains along the primary aminoacidic sequence (grey), and between non-consecutive subdomains (black). (d) Spring constants of intra-domain interactions between CG-sites, as a function of equilibrium lengths *x*_0_ for heteroENM D723R. (e) Spring constants for CG-sites of consecutive subdomains, versus *x*_0_ for heteroENM D723R. (f) Spring constants of CG-sites of non-consecutive subdomains, as a function of *x*_*0*_ for heteroENM D723R.

**Figure 6.**
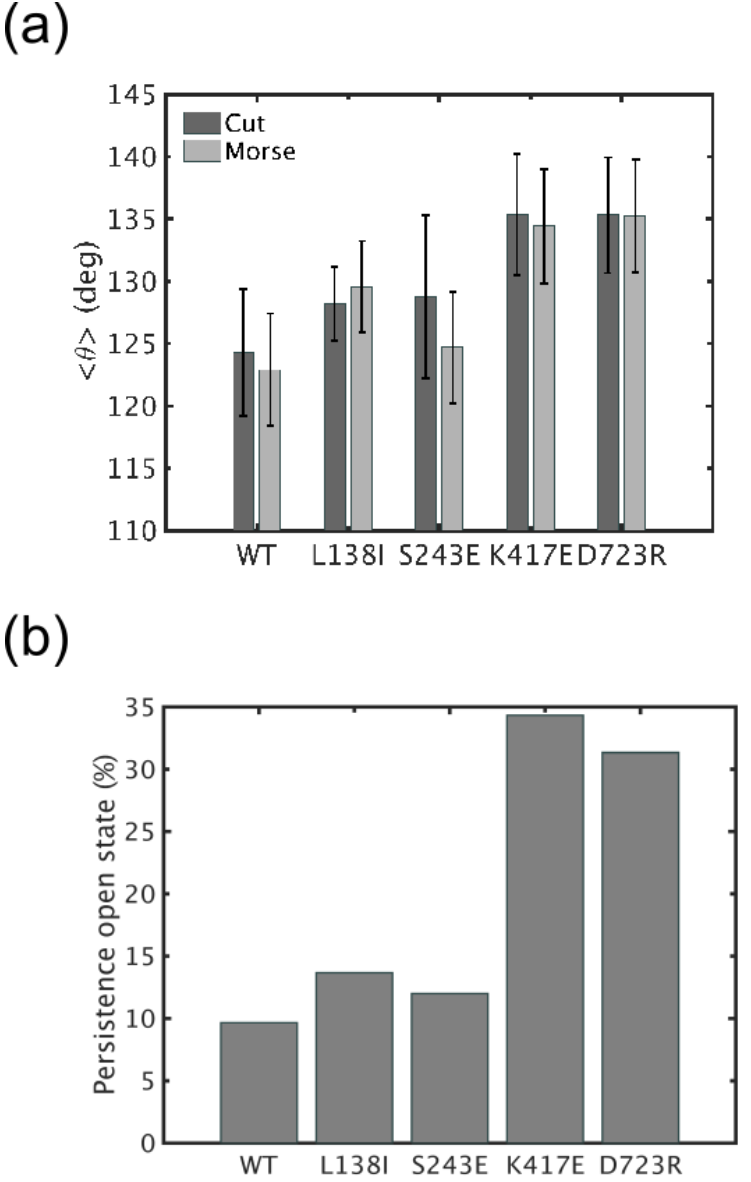
CG simulations of integrin mutants. (a) Average kink angle computed as mean of ϑ_1_ and ϑ_2_ over the course of the CG simulations, using WT and single mutant integrin with elastic springs above a threshold (*k* (*x*_0_) > 0.005 kcalmol^−1^A^2^) and either cut (eliminated) springs or soft Morse potentials below the threshold. (b) Persistence of the integrin open state, computed as the percentage of simulation time during which integrin has D_EM_ above 95%, its maximum value (results for cut springs above *k* (*x*_0_) > 0.1 kcalmol^−1^A^2^).

### Integrin affinity measurements

Next, we directly measured integrin affinity for the well-characterized Fab fragment, Wow-1, whose RGD-dependent binding to αvβ3 increases dramatically after activation (25). Wow1 is monomeric, minimizing possible effects of integrin clustering on its binding. Cells expressing WT or mutant αvβ3 were incubated with Wow-1 in standard, Mg^2+^ - and Ca^2+^ -containing buffer or in the presence of Mn^2+^, which is commonly used as a positive control for maximal activation (34). αvβ3-null cells were used as a negative control. We found that in Ca^2+^/Mg^2+^, all of the point mutants showed significantly increased binding compared to WT, with no significant differences detected among the activated mutants (Figure 7). Further, the activating mutants showed no further increase in the presence of Mn^2+^. This last result implies that all of the mutants are maximally activated in these assays.

**Figure 7.**
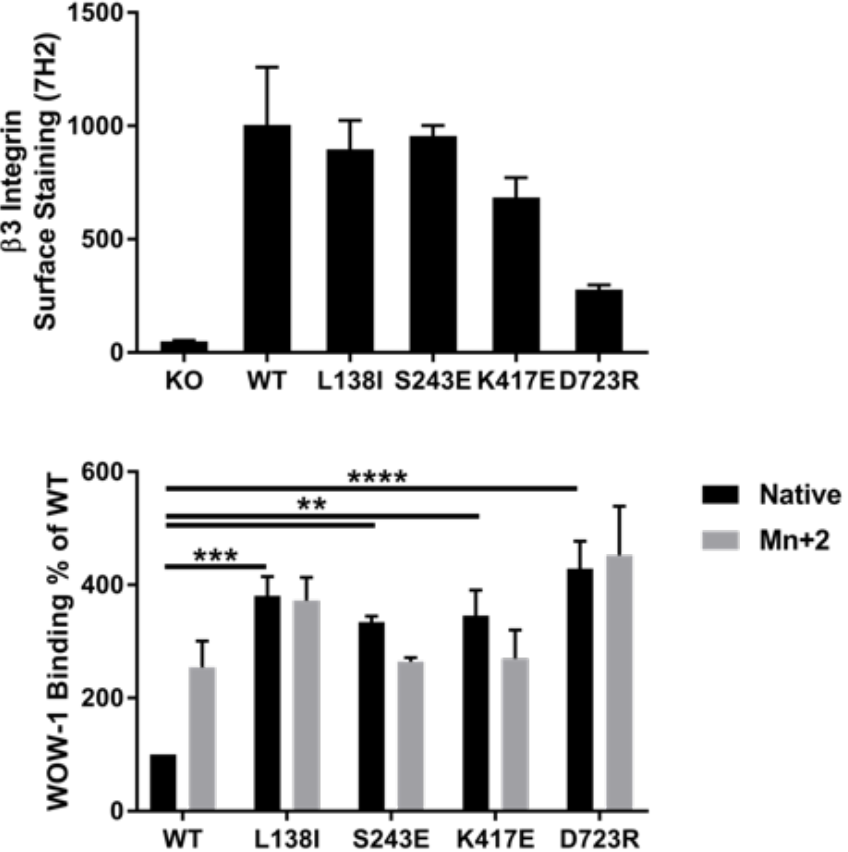
Integrin activation assays. α_v_β_3_ activation was assayed experimentally by binding of the monomeric, ligand-mimetic Fab, WOW-1. β3 integrin knockout cells expressing wild type or mutant β3 in suspension were stained Alexa647-conjugated antibody 7H2 that binds to all α_v_β_3_, and with Alexa488-conjugated WOW-1 by flow cytometry (~ 500k cells per condition). Binding of WOW-1 was normalized to the average total β3 integrin (a). Binding was performed under native, EDTA, and Mn^+2^ conditions. WOW-1 binding is shown relative to WT in native conditions (b). Values are means ± SEM, n=4 independent experiments. (** p < 0.01, *** p < 0.001, **** p<0.0001, Two-way ANOVA with Sidak’s multiple comparisons test)

### CG simulations reveal intermediate states

In all of our CG simulations, the integrin headpiece extended away from the legs, with the degree of extension depending on the mutant examined and the threshold of removed or converted harmonic interactions. Snapshots from a representative simulation of S243E, with springs preserved for k > 0.1 kcalmol^−1^A^2^, are shown in Figure 8. We next computed the RMSD of the simulated systems for all of the single point mutants relative to the four cryo-EM structures (24). In particular, we looked at the configurations of wild type and mutant integrins that were closest to each cryo-EM structure, using the minimum RMSD from CG trajectories, RMSDMIN, and compared how much they differed. Our results showed that for *k* > 0.001 kcalmol^−1^A^2^, the single mutants deviate from the cryo-EM conformer more than the WT in the following cases: closed (Figure 9a) and first intermediate (Figure 9b) and open conformer (Figure 9d) cryo-EM. With respect to the second intermediate from the cryo-EM conformer, the mutant systems generally have higher RMSD_MIN_ than the WT for *k* > 0.0001 kcalmol^−1^A^2^. The comparison of our CG mutants with full-length cryo-EM conformers from single molecule experiments show surprisingly strong structural similarity. This indicates that the simulations are consistent with a deadbolt model for integrin opening and inconsistent with the switchblade mechanism. Furthermore, the simulations show that this opening mechanism results from weakening low stiffness long-range correlations while preserving local dynamics correlations within each integrin subdomain and between pairs of neighboring subdomains along the primary protein sequence.

**Figure 8.**
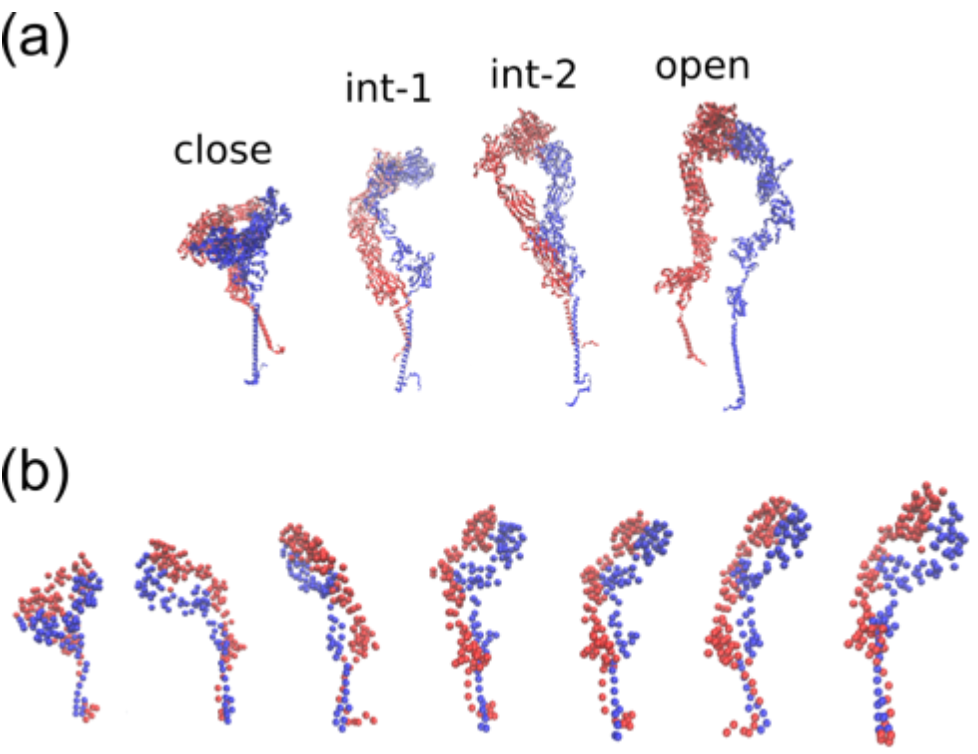
CG models show that headpiece extension occurs without legs separation. (a) reconstructed configurations form cryo-EM (24): closed; first intermediate; second intermediate and open conformers. (b) Representative snapshots from CG simulations of S243E with cut springs below *k*(*x*_0_) = 0.1 kcalmol^−1^A^2^, showing extension of the closed conformer.

**Figure 9.**
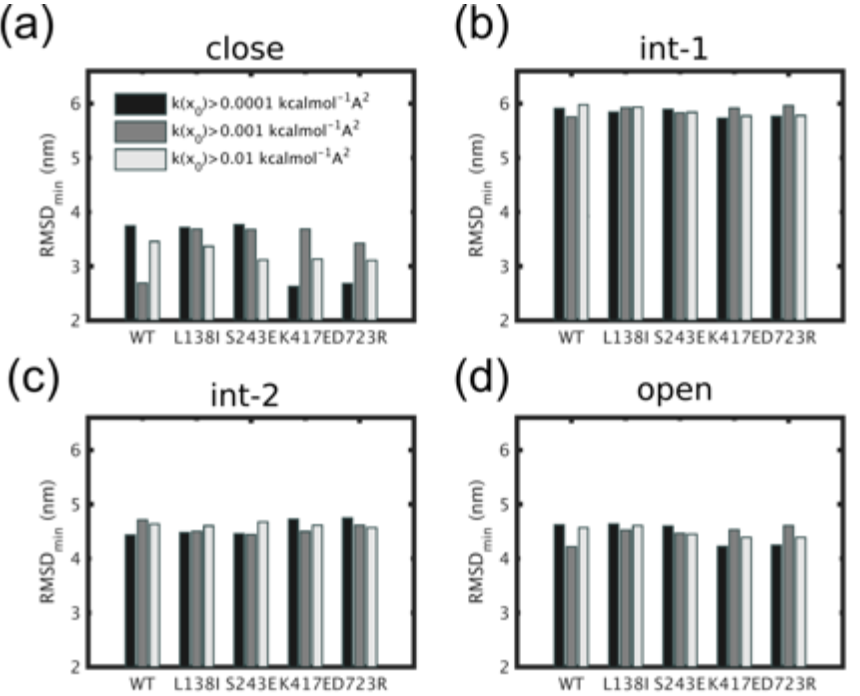
Comparison of CG-heteroENM structures with cryo-EM reconstructions of closed, intermediate and open integrins. (a) Minimum RMSD between CG-heteroENM simulations and the closed conformer. (b) Minimum RMSD between CG-heteroENM simulations and the first integrin intermediate. (c) Minimum RMSD between CG-heteroENM simulations and second integrin intermediate. (d) Minimum RMSD between CG-heteroENM simulations and open integrin conformer.

## DISCUSSION

In this study, we have addressed how integrin headpiece extension coordinates with leg separation during integrin opening. We utilized a novel combination of AA MD simulation, ED-CG modeling, and a modified heteroENM approach that allows for large conformational change on single and double point mutants of full-length α_V_β_3_ integrin. We investigated whether destabilization of the closed conformation occurs as localized structural rearrangements or as more cooperative changes in the receptor headpiece and legs. Also, we used ligand binding experiments and comparison of our CG models with single molecule cryoEM integrin reconstructions to validate our modeling configurations.

The AA MD simulations, albeit of limited large-scale sampling, showed that β3 mutations induce molecular structural rearrangements in both headpiece and legs of α and β subunits. In mutants, these rearrangements are generally enhanced relative to wild type integrins, by initiation of headpiece opening, flattening of the kink angle, and separation of the transmembrane helices (Figure 3). All mutants here analyzed destabilize the linkers within the α and β subunits, including Linker 1 and the EGF motifs (Figure 2b-c, e-f), respectively. The effect of mutants on linkers could facilitate structural transition towards open states. This result is in agreement with previous data from MD simulations on the EGF- motifs of the β subunit single cysteine mutations (35). These studies suggested that rearrangement of disulfide bonds in mutants could be part of a cascade of thiol/disulfide exchange reactions for activation (35). Results from our AA MD simulations also reveal that while structural properties of the systems do not converge within 1 μs (Figure S1 and Figure S3), the mutants that maximally destabilize the closed state are S243E and D723R (Figure S4). The hydrophilic surface area is known to be significantly larger for S243E compared to wild type αvβ3 and other mutants. This reflects greater extension of the headpiece away from the lipid bilayer, flattening of both kink angles and initiation of separation of transmembrane helices (Figure 3c and Figure S4). The effect of S243E on the global structural reorganization of integrin is triggered by the formation of a salt bridge at the point of the mutation, which induced local reorientation of a histidine, aspartic acid and arginine (Figure 4). In the case of the double mutant D723R_S243E, a salt bridge still forms in the βA domain (Figure S6), but the altered electrostatic interactions induced by mutation of aspartic acid to arginine in the β transmembrane helix and its higher pKa generate more stable interactions within the two helices (Figure S7).

The result that a single mutation in the βA domain of the β subunit can initiate structural opening is consistent with a number of previous MD studies of the integrin headpiece. For example, simulations of fibronectin-bound integrin headpiece showed that the ligand binding pocket at the interface between α and β subunits together with the hinge between the βA and hybrid domain of the β subunit are allosterically linked to initiate opening (18). In MD simulations of both α_IIb_β_3_ and α_V_β_3_ integrin headpieces, a common transition pathway for propagation of conformational changes within the βA domain was identified as the precursor of structural opening (36), consistent with our results from modified heteroENM that opening of the βA/hybrid junction can act as a hinge. Molecular simulations of the integrin headpiece were also previously performed in combination with experimental headpiece mutation to show that reorientation of the hybrid domain in the β subunit is required for structural activation (37). Forced unbending of the integrin αvβ3 headpiece was simulated using SMD, which showed that pulling the head readily induced changes starting from the headpiece (38). This supports that headpiece extension is very critical in integrin opening. In the current study, we used the full-length atomistic receptor which, unlike previous efforts, allowed us to characterize motional correlations between headpiece and legs and to further detect the impact of short versus long range correlations on integrin extension. With this study, we were also able to analyze the effect of the D723R mutation in the cytoplasmic tail of the β subunit, showing that propagation of structural activation can also be reproduced in this mutant.

In order to set our MD results into a broader context and test whether headpiece extension and leg separation are correlated at longer time scales, we used multiscale CG methods on individual integrins (with effect of lipid bilayer implicitly included in the AA MD input), based on ED-CG and HeteroENM methods, which were developed in (32, 33). This approach reduced the number of integrin sites from 27215 to 200, and reduced the computational cost by >100-fold. We modified our standard heteroENM model so that it can undergo large scale conformational changes by systematically removing low stiffness springs or converting them into softer, dissociable Morse interactions to facilitate realistic conformational flexibility. The predictive validity of standard ED-CG-hENM approaches was therefore significantly extended to enable sampling of multiple integrin conformations outside of the AA MD used to parameterize aspects of the model.

Results from the CG simulations confirmed that all mutants destabilized the integrin closed conformation via enhancement of kink angles and persistence of open configurations (Figure 6). This result was consistent with the experimental finding that WOW-1 binding was maximal for all of the mutants examined (Figure 7). However, this maximal activation for all mutants obscures possible differences. Dynamic measurements that are sensitive to the kinetics of activation will be an interesting target for future studies.

We also compared our αvβ3 CG structures with cryo-EM reconstructions of αIIbβ3 integrins at different states of activation, which had observed headpiece extension before leg separation (24). Integrin extension in the CG models resulted from preserving local atomistic dynamics correlations within each integrin subdomain and between pairs of neighboring subdomains along the primary protein sequence, while removing weak molecular long-range correlations. This suggests that, in the case of limited sampling, certain correlations present in the AA MD simulations were not representative of correlations in the global conformational landscape.

## CONCLUDING REMARKS

Our results from AA MD simulations show that coordinated atomistic motions within and between headpiece and legs destabilize integrin closed conformation on time scales on the order of μs (8). However, at longer time scales, accessed via CG simulations, headpiece extension precedes leg separation. Also, this activation mechanism for αvβ3 integrin is consistent with recent cryo-EM reconstructions of α_IIb_β_3_ integrin (20). In our simulations, extension of the headpiece occurs upon reduction of heteroENM effective harmonic interaction connectivity by maintaining connections within and between consecutive subdomains and modifying the low-frequency connections between distant subdomains. This implies that local contacts can persist during integrin opening and that long-range, low-frequency motional correlations are not consistent between closed and extended integrin states. Our model therefore supports the notion integrin extension results from disruption of weak, long-range interactions. In order for the legs to move apart in the model, stronger correlations between non-consecutive subdomains should also be reduced in the CG model, but this would lead to an overall loss of structural integrity and not only legs separation. Stated differently, our model is inconsistent with a switchblade mechanism.

To conclude, the main findings from this study are: (1) point mutations in the β subunit destabilize the αvβ3 closed structure in the absence of extracellular or intracellular ligands; (2) in the mutants, both integrin headpiece and legs respond to destabilization of the closed configuration via transmission of conformational transitions through flexible linkers; (3) the S243E mutant is an “activating” mutation that acts not only on the integrin headpiece but also allosterically on the transmembrane helices at the molecular level; and (4) headpiece extension can occur before leg separation, similar to cryo-EM reconstructions and consistent with a deadbolt model of integrin activation (Figure 8 and Figure 9).

Our findings of more general relevance are the following. Motions induced by integrin distal parts, that are weakly correlated at the atomistic level, leads to increased structural flexibility in full-length integrin and can thus promote extension. This motion can be induced experimentally in WT integrin by extracellular ligand binding or intracellular binding of the cytoskeletal protein talin, that bind the receptor in distal locations. Extracellular or intracellular integrin binding proteins can thus modify weakly correlated interactions between distal subdomains of the receptor, while preserving stronger, local correlations within subdomains and between neighboring subdomains. The idea of preserving collective local motions for integrin structural activation points towards a view of the receptor as sensitive in conformation to changes of weak, long-range inter-domains interactions.

## MATERIALS AND METHODS

In order to characterize integrin conformational activation, we computationally reconstructed full-length α_V_β_3_ integrins embedded in a lipid bilayer. We mutated the WT conformer using single or double point activating mutations in the βA domain of the β subunit, in the transmembrane β helix or in both. We tested whether initiation of integrin opening is, on the timescale of 1μs, a local effect, involving only the headpiece or the legs, or a global phenomenon, with simultaneous structural rearrangements in both the headpiece and the legs. Then, we built coarse-grained models of each integrin system to sample a wider range of conformations and test correlations between headpiece extension and leg separation over longer times. With both MD and CG simulations, we tested which activating mutations favor structural opening. This result was verified with experiments using the monovalent ligand WOW-1Fab (25), which direct assesses α_V_ß_3_ affinity state. Last, we compared our CG conformations with available cryo-EM reconstructions of α_IIb_β_3_ integrin, which is closely related to αvβ3 (24).

### All-atom simulations

To examine the effect of mutation on the conformation of full length integrin α_V_β_3_, μs length all–atom molecular dynamics (AA MD) simulations were performed on the integrins embedded in a lipid square patch (see Figure 1b). We assembled the bent headpiece of α_V_β_3_ integrin [from 3IJE.pdb (26)] with its transmembrane helical and cytoplasmic parts [taken from 2KNC.pdb (27)], using homology modeling to reconstruct missing residues (28). Point mutations were selected based on studies that identified mutants that increased affinity for RGD ligands (12). Starting from this initial configuration, we used the VMD software (29) to build five single mutants and four double mutants: D723R, L138I, E206T, S243E, K417E, D723R-L138I, D723R-E206T, D723R-S243E and D723R-K417E. For each integrin, we generated a multicomponent model lipid bilayer with 80% DOPC and 20% DOPS lipids, using CHARMM-GUI membrane builder (30). We then removed lipids in the center of the lipid patch to make a hole and inserted the integrin (Figure 1b). Last, we placed the membrane/integrin system within a rectangular box and filled its space with TIP3P water molecules and 150 mM NaCl, for a total of about 1.2 million atoms. In order to reorganize the lipids around the integrin, energy minimization was run for 5000 steps of steepest decent algorithm, followed by 50 ns of position restraint in a constant NPT ensemble, using the Berendsen thermostat. Production AA MD simulation were carried out, using Gromacs 5.0.4 (31), for 1μs in the NPT ensemble using Nose-Hoover thermostat and Parrinello-Rahman barostat to keep the temperature at 310 K and pressure at 1 atm. Long-range electrostatic interactions were incorporated through the PME method with a cut-off of 1 nm. The same cut-off value was used for Lennard-Jones interactions.

In order to identify differences in headpiece versus leg arrangements between WT integrin and the mutants, we defined metrics that report both headpiece extension and legs separation (Figure 3a). We quantified the following: kink angles, ϑ_1_ and ϑ_2_, for α and β subunits, respectively; their average, ϑ; transmembrane legs separation; D_12_, and headpiece extension, D_EM_ (see schematics in Figure 3A). The angle ϑ_1_ was defined in the α subunit as the angle between points A1 (center of mass of residues 82-85 in the β-propeller domain), A2 (center of mass of residues 599-602, between tight domain and Linker 1) and A3 (center of mass of restudies 963-966 in the α transmembrane helix); for the β subunit, ϑ_2_ was defined as the angle between points B1 (center of mass of residues 236-239 in the βA domain), B2 (center of mass of residues 480-484, between the hybrid domain and the EGF-1/EGF-2 motifs) and B3 (center of mass of restudies 696-699 in the β transmembrane helix); E is a point at the interface between the two headpiece subunits and was defined as the center of mass between points A1 and B1; M was defined as the center of mass between points A3 and B3 (corresponding residues are 963-966 and 696-699 in the two helices); points A and B, whose distance was indicated with D_12_, were given by the midpoints of each transmembrane helix.

### Coarse-grained model

In order to fully become active, integrins must sample multiple intermediate conformational states (24). Many of these conformational states were not accessed by AA MD simulations owing to the relatively short timescales that can be sampled at that level. We therefore built CG models based on the observed motional correlations of atoms in MD simulations and used them to identify structural differences between the WT and mutants on effectively longer timescales. We first developed Essential Dynamics Coarse-graining (ED-CG) (32) and heterogeneous elastic network (HeteroENM) models (33) of each integrin starting from the AA MD trajectories (without explicit inclusion of the lipid membrane in these model-it is there in the AA MD data, however). The ED-CG approach was chosen in order to select CG sites which preserve independent motion in the CG protein, and because it is constructed from the primary protein sequence without distorting it when exploring a wide conformational space. The heteroENM approach was used to create effective harmonic interactions between the CG sites which directly capture nanoscale correlations from the AA MD simulations. This approach was critical for creating CG models that maintain molecular differences between the integrin mutants studied here.

However, our present application of ED-CG and heteroENM differs from previous studies in one important and novel way. In previous applications of ED-CG and heteroENM, the elastic network was created to replicate all fluctuation dynamics observed in the reference AA MD simulations (32, 33). Here, certain correlations present in the MD simulations only provide a glimpse of the larger scale changes in the global conformational landscape, because of their limited sampling. In order for the CG model to sample substantially beyond the observed reference AA MD configurations, the hENM procedure was modified in this work by reducing the effective harmonic connectivity of each hENM integrin as a model for large conformation changes. We either systematically removed a varying fraction of the effective harmonic potentials or converted some of the inter-domain harmonic interactions into “softer” Morse potentials. In particular, we modified only those inter-domain interactions between non-consecutive subdomains along the subunits in order to maintain connections along the primary sequence of the proteins. We tested conditions where harmonic interaction potentials with equilibrium stiffness *k* < 0.0005-0.1 kcalmol^−1^A^2^ were modified, using 200 CG sites and an enforced cutoff 3 nm for each integrin. By modifying a fraction of harmonic potentials with *k* below 0.0005 kcalmol^−1^A^2^, no significant structural reconfiguration of the receptor was observed. By increasing the upper limit above 0.1 kcalmol^−1^A^2^, structural connectivity was lost. Upon performing CG dynamics on these systems, we evaluated their conformational motion in order to characterize whether destabilization of the closed integrin conformer occurs in a coordinated fashion between headpiece and legs, and detect which mutants most effectively destabilized the closed conformation. Last, we compared our CG trajectories with mapped cryo-EM reconstructions of α_IIb_β_3_ integrin intermediates (24).

### Cell Lines

Mouse lung endothelial cells (MLECs) were isolated from β3 integrin null mice. β3 integrin single point mutants were generously provided by Timothy Springer (Harvard University) and Mark Ginsberg (UCSD). Double mutants were constructed by standard site directed mutagenesis. These sequences were subcloned into pBOB vector and virus prepared in HEK 293Tx cells by co-transfecting with pCMV-VSV-G and psPAX2 using Lipofectamine 2000 (Invitrogen). The temperature sensitive mutant of the SV40 virus large T antigen was employed for conditional immortalization of these cells. Immortalized β3−/− MLECs were infected with wild-type (WT) or mutant β3 integrin viruses and subsequently sorted to obtain homogenous populations with equal expression levels. For expansion, MLECs were cultured in 1:1 Hams F-12 and low glucose DMEM with 20% FBS, 1% Penicillin-Streptomycin, 2.5mM glutamine and endothelial cell growth supplement (ECGS, 50mg/L) at the permissive temperature of 30°C. For experiments, cells were switched to 37°C to inactivate large T one day prior.

### Integrin activation measurements

To measure integrin affinity state, cells were detached using 0.25% trypsin, washed with complete medium, resuspended in serum-free medium at 13.3 × 10^6^ cells/mL, and incubated with primary β3-specific integrin ligands and secondary antibodies at 4 × 10^5^ cells in 50 μL as described (34). Briefly, to assess surface expression of β3, cells were incubated with 7H2 primary antibody (Developmental Studies Hybridoma Bank), washed with DMEM, incubated with anti-mouse-Alexafluor-647 secondary antibody (Invitrogen), washed with DMEM, and resuspended in PBS. To assess β3 activation state, cells were incubated with WOW-1 primary Fab (generously provided by S. Shattil) in the presence or absence of 20 mM EDTA or 2 mM MnCl_2_, washed with DMEM, incubated with secondary F(ab’)2 anti-mouse IgG (H+L) Alexafluor-488 (Invitrogen), washed with DMEM, and resuspended in PBS. Primary antibody was omitted to assess background fluorescence. Fluorescence was measured on an LSRII flow cytometer (BD Biosciences). The β3-integrin activation index of cells was calculated as AI = (F - Fo)/Fintegrin, where F is the geometric mean fluorescence intensity (MFI) of WOW-1 binding after background subtraction, Fo is the MFI of WOW-1 binding in presence of EDTA inhibitor, and Fintegrin is the normalized MFI of 7H2 antibody binding to cells. Activation was normalized to WT under native conditions for 4 independent experiments.

## ACKNOWLEDGEMENTS

This research was supported by the DOD/ARO through a MURI grant W911NF1410403. This work was also partially supported by the University of Chicago Materials Research Science and Engineering Center, which is funded by National Science Foundation under award number DMR-1420709. Computational time was provided by the National Science Foundation XSEDE resources at the Pittsburgh Supercomputing Center.

## REFERENCES

1. Hood, J.D., and D.A. Cheresh. 2002. ROLE OF INTEGRINS IN CELL INVASION AND MIGRATION. Nat. Rev. Cancer. 2: 91–100.

2. Bershadsky, A.D., N.Q. Balaban, and B. Geiger. 2003. Adhesion-Dependent Cell Mechanosensitivity. Annu. Rev. Cell Dev. Biol. 19: 677–695.

3. Ingber, D.E. 2003. Mechanobiology and diseases of mechanotransduction. Ann Med. 35: 1–14.

4. Ross, T.D., B.G. Coon, S. Yun, N. Baeyens, K. Tanaka, M. Ouyang, and M.A. Schwartz. 2013. Integrins in mechanotransduction. Curr. Opin. Cell Biol. 25: 613–8.

5. Vogel, V., and M. Sheetz. 2006. Local force and geometry sensing regulate cell functions. Nat. Rev. Mol. Cell Biol. 7: 265–275.

6. Banno, A., and M.H. Ginsberg. 2008. Integrin activation. Biochem. Soc. Trans. 36: 229–234.

7. Carman, C. V, and T.A. Springer. 2003. Integrin avidity regulation: are changes in affinity and conformation underemphasized? Curr. Opin. Cell Biol. 15: 547–56.

8. Luo, B.-H., C. V Carman, and T.A. Springer. 2007. Structural basis of integrin regulation and signaling. Annu. Rev. Immunol. 25: 619–47.

9. Ye, F., C. Kim, and M.H. Ginsberg. 2012. Reconstruction of integrin activation. Blood. 119: 26–33.

10. Hynes, R.O. 2002. Integrins: bidirectional, allosteric signaling machines. Cell. 110: 673–87.

11. Campbell, I.D., and M.J. Humphries. 2011. Integrin Structure, Activation, and Interactions. Cold Spring Harb. Perspect. Biol. 3: a004994–a004994.

12. Luo, B.-H., J. Karanicolas, L.D. Harmacek, D. Baker, and T.A. Springer. 2009. Rationally designed integrin beta3 mutants stabilized in the high affinity conformation. J. Biol. Chem. 284: 3917–24.

13. Luo, B.-H., T.A. Springer, and J. Takagi. 2003. Stabilizing the open conformation of the integrin headpiece with a glycan wedge increases affinity for ligand. Proc. Natl. Acad. Sci. U. S. A. 100: 2403–8.

14. Luo, B.-H., K. Strokovich, T. Walz, T.A. Springer, and J. Takagi. 2004. Allosteric _NL_ 1 Integrin Antibodies That Stabilize the Low Affinity State by Preventing the Swing-out of the Hybrid Domain*.

15. Xie, C., J. Zhu, X. Chen, L. Mi, N. Nishida, and T.A. Springer. 2010. Structure of an integrin with an αI domain, complement receptor type 4. EMBO J. 29: 666–679.

16. Xiao, T., J. Takagi, B.S. Coller, J.-H. Wang, and T.A. Springer. 2004. Structural basis for allostery in integrins and binding to fibrinogen-mimetic therapeutics. Nature. 432: 59–67.

17. Huth, J.R., E.T. Olejniczak, R. Mendoza, H. Liang, E.A. Harris, M.L. Lupher, A.E. Wilson, S.W. Fesik, D.E. Staunton, and D.E. Staunton. 2000. NMR and mutagenesis evidence for an I domain allosteric site that regulates lymphocyte function-associated antigen 1 ligand binding. Proc. Natl. Acad. Sci. U. S. A. 97: 5231–6.

18. Puklin-Faucher, E., M. Gao, K. Schulten, and V. Vogel. 2006. How the headpiece hinge angle is opened: new insights into the dynamics of integrin activation. J. Cell Biol. 175: 349–360.

19. Mehrbod, M., S. Trisno, and M.R.K. Mofrad. 2013. On the activation of integrin αIIbβ3: outside-in and inside-out pathways. Biophys. J. 105: 1304–15.

20. Adair, B.D., J.-P. Xiong, C. Maddock, S.L. Goodman, M.A. Arnaout, and M. Yeager. 2005. Three-dimensional EM structure of the ectodomain of integrin {alpha}V{beta}3 in a complex with fibronectin. J. Cell Biol. 168: 1109–18.

21. Mould, A.P., S.J. Barton, J.A. Askari, P.A. McEwan, P.A. Buckley, S.E. Craig, and M.J. Humphries. 2003. Conformational changes in the integrin beta A domain provide a mechanism for signal transduction via hybrid domain movement. J. Biol. Chem. 278: 17028–35.

22. Zhu, J., J. Zhu, and T.A. Springer. 2013. Complete integrin headpiece opening in eight steps. J. Cell Biol. 201.

23. Arnaout, M.A., S.L. Goodman, and J.-P. Xiong. 2007. Structure and mechanics of integrin-based cell adhesion. Curr. Opin. Cell Biol. 19: 495–507.

24. Xu, X.-P., E. Kim, M. Swift, J.W. Smith, N. Volkmann, and D. Hanein. 2016. Three-Dimensional Structures of Full-Length, Membrane-Embedded Human α(IIb)β(3) Integrin Complexes. Biophys. J. 110: 798–809.

25. Pampori, N., T. Hato, D.G. Stupack, S. Aidoudi, D.A. Cheresh, G.R. Nemerow, and S.J. Shattil. 1999. Mechanisms and consequences of affinity modulation of integrin alpha(V)beta(3) detected with a novel patch-engineered monovalent ligand. J. Biol. Chem. 274: 21609–16.

26. Xiong, J.-P., B. Mahalingham, J.L. Alonso, L.A. Borrelli, X. Rui, S. Anand, B.T. Hyman, T. Rysiok, D. Müller-Pompalla, S.L. Goodman, and M.A. Arnaout. 2009. Crystal structure of the complete integrin αVβ3 ectodomain plus an α/β transmembrane fragment. J. Cell Biol. 186: 589–600.

27. Yang, J., Y.-Q. Ma, R.C. Page, S. Misra, E.F. Plow, and J. Qin. 2009. Structure of an integrin alphaIIb beta3 transmembrane-cytoplasmic heterocomplex provides insight into integrin activation. Proc. Natl. Acad. Sci. U. S. A. 106: 17729–34.

28. Pettersen, E.F., T.D. Goddard, C.C. Huang, G.S. Couch, D.M. Greenblatt, E.C. Meng, and T.E. Ferrin. 2004. UCSF Chimera?A visualization system for exploratory research and analysis. J. Comput. Chem. 25: 1605–1612.

29. Humphrey, W., A. Dalke, and K. Schulten. 1996. VMD: Visual molecular dynamics. J. Mol. Graph. 14: 33–38.

30. Jo, S., J.B. Lim, J.B. Klauda, and W. Im. 2009. CHARMM-GUI Membrane Builder for Mixed Bilayers and Its Application to Yeast Membranes. Biophys. J. 97: 50–58.

31. Abraham, M.J., T. Murtola, R. Schulz, S. Páll, J.C. Smith, B. Hess, and E. Lindah. 2015. Gromacs: High performance molecular simulations through multi-level parallelism from laptops to supercomputers. SoftwareX. 1-2: 19–25.

32. Zhang, Z., L. Lu, W.G. Noid, V. Krishna, J. Pfaendtner, and G.A. Voth. 2008. A Systematic Methodology for Defining Coarse-Grained Sites in Large Biomolecules. Biophys. J. 95: 5073–5083.

33. Lyman, E., J. Pfaendtner, and G.A. Voth. 2008. Systematic multiscale parameterization of heterogeneous elastic network models of proteins. Biophys. J. 95: 4183–92.

34. Bouaouina, M., D.S. Harburger, and D.A. Calderwood. 2011. Talin and Signaling Through Integrins. In: Methods in molecular biology (Clifton, N.J.). . pp. 325–347.

35. Levin, L., E. Zelzion, E. Nachliel, M. Gutman, Y. Tsfadia, and Y. Einav. 2013. A Single Disulfide Bond Disruption in the β3 Integrin Subunit Promotes Thiol/Disulfide Exchange, a Molecular Dynamics Study. PLoS One. 8: e59175.

36. Provasi, D., M. Murcia, B.S. Coller, and M. Filizola. 2009. Targeted molecular dynamics reveals overall common conformational changes upon hybrid domain swing-out in β3 integrins. Proteins Struct. Funct. Bioinforma. 77: 477–489.

37. Cheng, M., J. Li, A. Negri, B.S. Coller, and L.R. Languino. 2013. Swing-Out of the b3 Hybrid Domain Is Required for aIIbb3 Priming and Normal Cytoskeletal Reorganization, but Not Adhesion to Immobilized Fibrinogen.

38. Chen, W., J. Lou, J. Hsin, K. Schulten, S.C. Harvey, and C. Zhu. 2011. Molecular Dynamics Simulations of Forced Unbending of Integrin αVβ3. PLoS Comput. Biol. 7: e1001086.

